# High-resolution mass spectrometry reveals environmentally relevant uptake, elimination, and metabolic alterations following early embryonic exposure to 2,3,7,8-tetrachlorodibenzo-p-dioxin in zebrafish

**DOI:** 10.1101/2022.05.04.490602

**Authors:** Michelle E. Kossack, Katherine E. Manz, Nathan R. Martin, Kurt D. Pennell, Jessica Plavicki

## Abstract

Dioxin and dioxin-like compounds are ubiquitous environmental contaminants that induce toxicity by binding to the aryl hydrocarbon receptor (AHR), a ligand activated transcription factor. The zebrafish model has been used to define the developmental toxicity observed following exposure to exogenous AHR ligands such as the potent agonist 2,3,7,8-tetrachlorodibenzo-p-dioxin (dioxin, TCDD). While the model has successfully identified cellular targets of TCDD and molecular mechanisms mediating TCDD-induced phenotypes, fundamental information such as the body burden produced by standard exposure paradigms is still unknown. We performed targeted gas chromatography (GC) high-resolution mass spectrometry (HRMS) in tandem with non-targeted liquid chromatography (LC) HRMS to quantify TCDD uptake, model the elimination dynamics of TCDD, and determine how TCDD exposure affects the zebrafish metabolome. We found that 10 ppb, 1 ppb, and 50 ppt waterborne exposures during early embryogenesis produced environmentally relevant body burden of TCDD: 38 ± 4.34, 26.6 ± 1.2, and 8.53 ± 0.341 pg/embryo, respectively, at 24 hours post fertilization. In addition, we discovered that TCDD exposure was associated with the dysregulation of several metabolic pathways that are critical for brain development and function including glutamate metabolism, chondroitin sulfate biosynthesis, and tyrosine metabolism pathways. Together, these data demonstrate that existing exposure paradigms produce environmentally relevant body burdens of TCDD in zebrafish and provide insight into the biochemical pathways impacted by toxicant-induced AHR activation.

**HIGHLIGHTS:**

- Historical TCDD exposure paradigms produce environmentally relevant body burdens in zebrafish embryos.
- TCDD elimination for high doses can be modeled using an exponential regression.
- Exposure to TCDD alters metabolic pathways that are essential for brain development and function.

**GRAPHICAL ABSTRACT:** 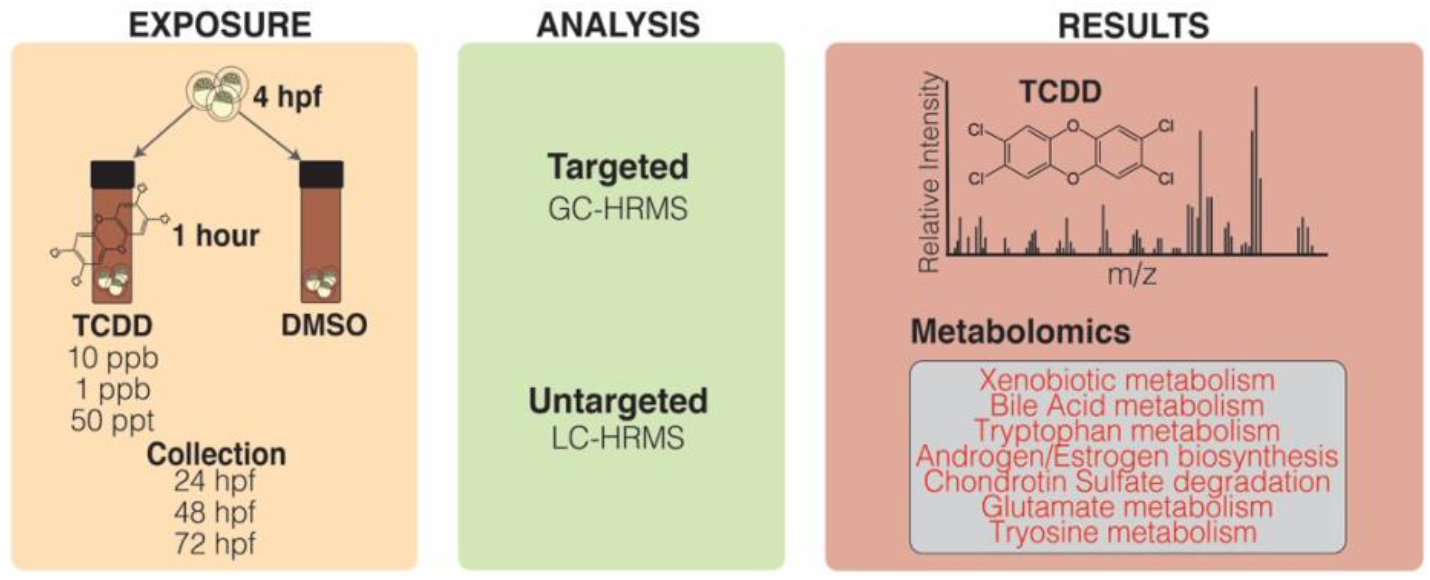

## INTRODUCTION

Dioxin and dioxin-like compounds are global contaminants that pose a threat to human health as carcinogens, immunomodulators, neurotoxicants, endocrine disruptors, and teratogens (1). While some dioxins are produced through natural processes such as volcanic activity and forest fires, the majority of dioxin and dioxin-like compounds are introduced into the environment through anthropomorphic activities such as paper production and the burning of industrial waste (2). Dioxins induce toxicity by binding the aryl hydrocarbon receptor (AHR), a ligand activated transcription factor with endogenous functions in vascular, brain, and immune system development and function (3–13). Hundreds of dioxins and dioxin-like congeners exist in the environment with each congener having varying affinity to AHR (14–16). The potent AHR agonist, 2,3,7,8-tetrachlorodibenzo-p-dioxin (TCDD), is routinely used in laboratory models to study developmental toxicity induced by exogenous AHR ligands.

Zebrafish have become a popular model system in developmental biology and toxicology due to their rapid external embryonic development, optical transparency, and high degree of sequence homology to humans (17). In zebrafish, embryonic exposure to TCDD for 1 hour at a concentration of either 1 parts-per-billion (ppb) or 10 ppb is sufficient for sustained AHR activation and produces developmental defects including cardiovascular malformations, hemorrhaging, loss of the proepicardial progenitor cells, cardiac and cerebral edema, and craniofacial defects (18–24). Studies in zebrafish have revealed that one of the critical molecular mechanisms underlying a number of the observed TCDD-induced phenotypes is the reduced expression and function of the high mobility group transcription factor *sox9b* (22,25,26). TCDD-induced inhibition of *sox9b* in developing zebrafish is mediated by the induction of *sox9b long intergenic noncoding RNA* (*slincR*) (27,28). These mechanistic findings derived from model organism studies make important contributions to our understanding of how human exposure to TCDD may disrupt human development and health. In humans, dysregulation of *SOX9*, the human paralog of *sox9b*, is known to cause skeletal, central nervous system, heart malformations, phenotypes which mirror those observed in model organisms exposed to TCDD (29–33). The proposed human ortholog of *slincR*, LINC00673, was identified in the human genome, allowing for the possibility that a conserved mechanism mediates TCDD-induced toxicity in both zebrafish and humans (27). Although further research is needed, scientists continue to successfully use the zebrafish model to identify the molecular mechanisms underlying the teratogenic, carcinogenic, and other adverse health endpoints seen in humans following AHR agonist exposure.

While the zebrafish model has been used successfully to identify the cellular targets of TCDD and to determine the molecular mechanisms mediating toxicant induced phenotypes; the uptake and elimination dynamics of TCDD in zebrafish are not known for doses of TCDD that are commonly used in developmental exposure paradigms (34). TCDD is highly lipophilic with low solubility in water (K_ow_=7.02, (35)), which creates many questions relating to waterborne exposure paradigms, such as: How many grams of TCDD are present in the embryo following a waterborne exposure given the limited solubility of TCDD in water and its partitioning to other surfaces during the exposure? How quickly is TCDD eliminated from the developing embryo? What concentration of toxicant is bioavailable to the embryos? Does exposure to a high, verses a low, dose change uptake and elimination? Without this fundamental information, we are unable to interpret the environmental relevance of current dosing paradigms and whether the zebrafish exposure model reflects levels of dioxin exposure likely to occur in different human populations. To address this critical data gap, we used gas chromatography (GC) high-resolution mass spectrometry (HRMS) to quantify TCDD absorption in zebrafish embryos and non-targeted liquid chromatography (LC) HRMS metabolomics to understand how TCDD exposure during early embryogenesis affects the metabolome. We performed a one-hour waterborne TCDD exposures at 4 hours post fertilization (hpf) with three different concentrations of TCDD that are routinely used in zebrafish TCDD exposure studies: 50 parts-per-trillion (ppt), 1 part-per-billion (ppb), and 10 ppb. The 50 ppt exposure is used to model sublethal exposures and has been applied in transgenerational TCDD exposure studies, whereas the 1 ppb and 10 ppb exposures produce lethal cardiac phenotypes (23,36,37). Our metabolomic analysis revealed that TCDD exposure alters xenobiotic metabolism, tryptophan metabolism, and androgen biosynthesis. Based on knowledge derived from previous toxicological and biological studies these pathways were expected and detection was used to authenticate our metabolomic analysis. In addition, we discovered TCDD exposure was associated with the dysregulation of several metabolic pathways that are critical for brain development and function including the glutamate metabolism, chondroitin sulfate biosynthesis, and tyrosine metabolism pathways. Our findings provide essential data regarding TCDD uptake and elimination that are critical for future interpretation of studies using the zebrafish model to understand TCDD-induced toxicity. Furthermore, our metabolomics data provide new insight into the pathophysiological changes that contribute to phenotypes observed following TCDD exposure and provide new research directions for scientists to explore.

## MATERIALS AND METHODS

### Chemicals

TCDD stock solution for zebrafish exposure was purchased at an initial concentration of 50.00 ± 0.32 μg/mL (Stock A) in dimethyl sulfoxide (DMSO, ED-901-B, Cerriliant, St. Louis, MO). The stocks were made in 2 mL amber vials and caps were wrapped with Parafilm^®^. Dosing stocks were kept in the dark at room temperature. Dioxin mix, which contains TCDD, certified reference standard was purchased from AccuStandard for TCDD quantitation (New Haven, CT, USA). Surrogate standards (PCB 65 and 166) and internal standard mix (Phenanthrene-D10, Chrysene-D12, and Carbon Number Distribution Marker) were also purchased from AccuStandard. Solvents, including n-hexane (≥99%), acetone (99.8%, HPLC grade), dichloromethane (99.8%, HPLC grade), water (UHPLC-MS grade), and acetonitrile (UHPLC-MS grade) were purchased from Fisher Scientific. Isotopically labeled metabolites were purchased from Cambridge Isotopes (MSK-QC-KIT, Tewksbury, MA).

### Zebrafish Spawning and Exposure

All procedures involving zebrafish were approved by the Institutional Animal Care and Use Committee (IACUC) at the Brown University and adhered to the National Institute of Health’s “Guide for the Care and Use of Laboratory Animals”. AB strain zebrafish (*Danio rerio*) were maintained using a zebrafish aquatic housing system with centralized filtration, temperature control (28.5 ± 2°C), illumination (14 h:10 h light-dark cycle) ultraviolet (UV) germicidal irradiation, reverse osmosis (RO) water purification, and pH and conductivity monitoring (Aquaneering Inc., San Diego, CA) according to Westerfield (38). The Plavicki Lab Zebrafish Facility undergoes routine monitoring for disease including the semiannual qPCR panels to detect common fish pathogens.

Adult AB zebrafish were incrossed for 1 hour in 1.7 L slope breeding tanks (Techniplast, USA) and fertilized embryos were collected in egg water (1.5 mL stock salts in 1 L RO water). Embryonic and larval zebrafish were maintained at 28.5 ± 1°C in an incubator (Powers Scientific Inc., Pipersville, PA) within 100 mm non-treated culture petri dishes (CytoOne, Cat. No. CC7672-3394). Embryos were screened for fertilization and proper development prior to toxicant exposure.

When the embryos reached 4 hpf, 20 embryos were placed into a clean 2 mL amber vial (Cat. No. 5182-0558, Agilent, Santa Clara, CA) and residual egg water was removed. Fresh egg water (2 mL) was added to the vial (10 embryos/mL egg water) and then 2μL (1 μL TCDD/mL egg water) of TCDD stock (10,000 ng/mL or 50 ng/mL) or control Dimethyl Sulfoxide (DMSO) was added to the treatment to achieve a 10 ppb (10 ng/mL) or 50 ppt (0.05 pg/mL) final exposure concentration, respectively. Immediately after addition of TCDD, the vials were capped, wrapped in Parafilm^®^, and inverted several times to mix. To minimize inter sample variability, the same person performed all TCDD exposures. Treatment vials were placed on a rocker table for 1 hour at room temperature. Treatment solution was removed from the vials and embryos were washed three times with fresh egg water. Embryos were transferred to 6-well plates (20 embryos/well) containing egg water and placed into the incubator for collection.

### Collection and fixing

At 24 hpf, all embryos were manually dechorionated. Exposed and control embryos were fixed at either 24, 48, or 72 hpf. At 10 ppb, seven embryos were pooled per sample. At 50 ppt, 20-40 embryos were pooled per sample. All dosed embryo samples were paired with equivalent control samples. The water was removed from each sample before snap freezing in liquid nitrogen and storing at −80°C until ready for TCDD quantification and metabolomics analysis. At each timepoint 2-3 replicates were collected. In total there were 9 independent spawning events, each timepoint and dose was represented by at least three independent experiments totaling 5-7 measured samples per timepoint.

### Quantification of TCDD

Embryos were extracted and measured for TCDD concentrations by GC-HRMS (39). Briefly, pooled embryos were transferred into an amber glass 4-dram vial with 4 mL 1:1:1 hexane:acetone:dichloromethane. Each sample was spiked with 10 μL of surrogate standard solution containing PCB 65 and 166, to assess extraction recovery. The glass vial containing the embryos was sonicated for 2 hours, until the embryos were disintegrated, and mixed overnight (15 hours) on an orbital shaker. The supernatant was transferred to a QuEChERS (Quick, Easy, Cheap, Effective, Rugged, Safe) extraction tube containing 150 mg dispersive C18 powder and 900 mg MgSO_4_ (United Chemical Technologies, Bristol, PA, USA). The QuEChERS tube was mixed for 15 minutes and centrifuged for 10 minutes at 5,000 rpm. The supernatant from the QuEChERS tube was transferred to a glass test tube for collection. The 4-dram vial containing the embryos was rinsed twice with 1.5 mL of hexane:acetone:dichloromethane. Each rinse was transferred to the QuEChERS tube, mixed for 15 minutes in the QuEChERS tube, centrifuged, and transferred to the collection glass test tube. The final extract (7 mL total) was evaporated to 0.5-1 mL at 40°C under nitrogen using a 30 position Multivap Nitrogen Evaporator (Organomation Associates Inc.), transferred to a clear GC vial, and reduced to a final volume of 150 μL. The final extract was transferred to an amber autosampler vial containing a 250 μL glass insert, spiked with 10 μL of an internal standard solution containing 62.5 ug/L of phenanthrene D-10, chrysene D-12, and Carbon Number Distribution Marker and sealed with a Teflon^®^-lined screw cap.

Sample extracts were analyzed for TCDD concentration on a Thermo Q Exactive Orbitrap equipped with a Thermo Trace 1300 gas chromatograph (GC) and TriPlus RSH autosampler. 4 μL of the sample extracted were injected onto a 290°C split/splitless inlet operated in splitless mode. Sample extracts were separated on a Restek Rxi-35Sil MS (30 m x 0.25 mm inner diameter x 0.25 μm film thickness) column with helium (99.9999% purity) as the carrier gas at a flow rate of 1 mL/minute. The oven temperature started at 50°C for 0.5 minutes, increased to 330°C at 12°C/minute, and held at 330°C for 5 minutes (total run time was 28 minutes). The transfer line was maintained at 310°C. The source was held at 250°C and operated in electron ionization (EI) mode. Data were collected in full-scan mode (30-550 m/z). TCDD was quantified in TraceFinder 5.0 using the extracted ion chromatogram (XIC) and the most abundant peak in the mass spectrum (321.8928 m/z). TCDD identity was confirmed using the ratio of two confirming ions (319.8958 and 256.9321 m/z) and retention time (19.77 minutes). An eight-point calibration curve prepared by serial dilution of calibration standards in hexane (0.025 to 15 μg/L) was used to quantify. The limit of detection (LOD) was determined from seven injections of a calibration standard and calculated as: LOD = [*s* * *t* (*df*, 1 – α = 0.99)] / *m* where *s* is the standard deviation, t is the student’s t-value, *df* is the degrees freedom, α is the significance level (n=7, α =0.01, t=3.14), and *m* is the slope of the calibration curve (40). The LOD for TCDD was 0.0717 ng/mL (equivalent to 1.54 x 10^-6^ μg/embryo for 7 embryos or 2.69 x 10^-7^ μg/embryo for 40 embryos).

### High resolution metabolomics

Embryo samples were evaporated to dryness under nitrogen reconstituted in 150 μL acetonitrile containing a mixture of isotopically labelled metabolites. Non-targeted analysis for metabolite detection was performed by injecting 10 μL of sample extract on a Thermo Orbitrap Q Exactive HF-X MS equipped with a Thermo Vanquish ultra-high performance liquid chromatograph (UHPLC) system in triplicate. Two chromatography separation methods were used, normal and reverse-phase. The normal-phase LC was performed with a HILIC column (Thermo Syncronis HILIC 50 mm X 2.1 mm x 3 μm) at a constant temperature of 25°C. Mobile phase A contained 2 mM ammonium acetate in acetonitrile and mobile phase B contained 2 mM aqueous ammonium acetate. Metabolites were eluted from the column at a constant flow rate of 0.2 mL/minute using a solvent gradient as follows: equilibrate with 10% B for 1 minute, increase to 65% B for 9 minutes and hold for 3 minutes, decrease to 10% over 1 minute and hold for 1 minute. The reverse-phase LC was performed with a C18 column (Thermo Hypersil Gold Vanquish, 50 mm X 2.1 mm x 1.9 μm) at a constant temperature of 60°C. Mobile phase A contained 2 mM aqueous ammonium acetate and mobile phase B contained 2 mM ammonium acetate in acetonitrile. Metabolites were eluted from the column at a constant flow rate of 0.5 mL/minute using a mobile phase gradient as follows: equilibration with 2.5% B for 1 minute, increase to 100% B over 11 minutes and held for 2 minutes, and back to 2.5% B over 1 minute and held for 1.5 minutes (total run time 16.5 minutes, data were collected from 0.05 to 12.5 minutes). For both normal and reverse-phase LC, the MS was operated in full scan mode with 120,000 resolution, automatic gain control of 3 x 10^6^, and maximum dwell time of 100 ms. Electrospray ionization was conducted in positive mode for normal-phase and negative mode for reverse phase LC. Ionization was performed at a sheath gas flow of 40 units, auxiliary gas flow of 10 units, sweep gas flow of 2 units, spray voltage of 3.5 kV, 310°C capillary temperature, funnel radio frequency (RF) level of 35, and 320°C auxiliary gas heater temperature.

### Metabolomics data analysis

Data files were converted from *.raw files to *.cdf files using XCalibur file Converter, and then processed in R packages apLCMS (16) and xMSanalyzer (41) to produce m/z feature tables. The m/z feature tables were batch corrected using ComBat (42). Intensities were filtered by comparing peak intensity in the egg water blank to the sample intensity (intensity in the sample ≥ 1.5 times the blank) and normalizing by log2 transformation. Association of metabolite features obtained for 50 ppt and 10 ppb TCDD exposure levels was assessed for each exposure time and dose using t-tests (p < 0.05) in comparison to the control samples. A False Discovery Rate (FDR) ≤ 20% was used to control for Type I errors in multiple comparisons. Significant metabolites were analyzed for pathway enrichment using MetaboAnalystR (43) using the zebrafish mummichog curated model, which includes the KEGG, BiGG, and Edinburgh maps. All metabolomics data analysis was performed in R (version 4.0.2).

## RESULTS AND DISCUSSION

### TCDD body burdens following waterborne 10 ppb or 50 ppt exposures

The developmental toxicology community faces a number of historical issues that contribute to experimental variability and can make it difficult to interpret the environmental relevance of exposure paradigms. These challenges include the lack of standardized dosing paradigms, and the failure to implement quality control standards in research studies such as the validation of toxicant dosing solutions and the quantification of internal body burdens post-toxicant exposure. In light of these issues, we conducted a series of studies to determine TCDD uptake and elimination for dosing concentrations commonly used in TCDD exposure studies.

First, we diluted our commercially purchased and validated TCDD stock solution (stock A, Figure S1) in DMSO to make a 10,000 ppb secondary stock solution (Stock B), which was serial diluted to make a 1,000 ppb stock (Stock C) and 50 ppb stock solution (50,000 ppt, Stock D). We validated the concentrations of each stock solutions (Figure S1), and found each stock was within the margin of error expected based on the purchased solution (Stock A). Each stock solution is diluted 1:1000 in embryo water to create a “working solution” to which the embryos are exposed. To validate that, not only our dilution was precise but also consistent between doses, we measured the TCDD concentration in three independent working solutions. We found that the working solutions were within expected deviation of the expected concentration (Figure S1). After validating all of the stock and working solutions, we were confident the stocks could be used for subsequent TCDD exposure and uptake studies.

For our body burden assessments, embryos were dosed with 50 ppt, 1 ppb, or 10 ppb TCDD at 4 hours post-fertilization (hpf) for a one-hour and then raised in fresh egg water. We selected the 1 ppb and 10 ppb doses, because these concentrations have been previously used to model TCDD-induced developmental toxicity and to identify key molecular mechanisms mediating TCDD-induced teratogenicity (23,25,27,37,44,45). The 50 ppt dose has been established as a sublethal concentration in embryonic and juvenile zebrafish (36,46–48) and used to model transgenerational effects of TCDD exposure. Changes in cardiac dysfunction have been reported as early as 60 hpf in the 1 and 10 ppb TCDD-exposed embryos; however, cardiac collapse and overt morphological malformations occur later in development and culminate in death between 5- and 7-days post fertilization (dpf, 20,37,47–49). Therefore, to avoid the confounds associated with organ dysfunction and overt morphological malformations, we selected 24, 48, and 72 hpf as our time points of interest for both the body burden and metabolomic studies (Figure 1). Consistent with previous reports, we did not observe differences in mortality between the control and exposure group at any time point. The average survival of larvae at 72 hpf in the control (DMSO) group was 96.4% and 96.3% in the TCDD group (p=0.49, one-tailed t-test, n=313 fish).

**Figure 1.**
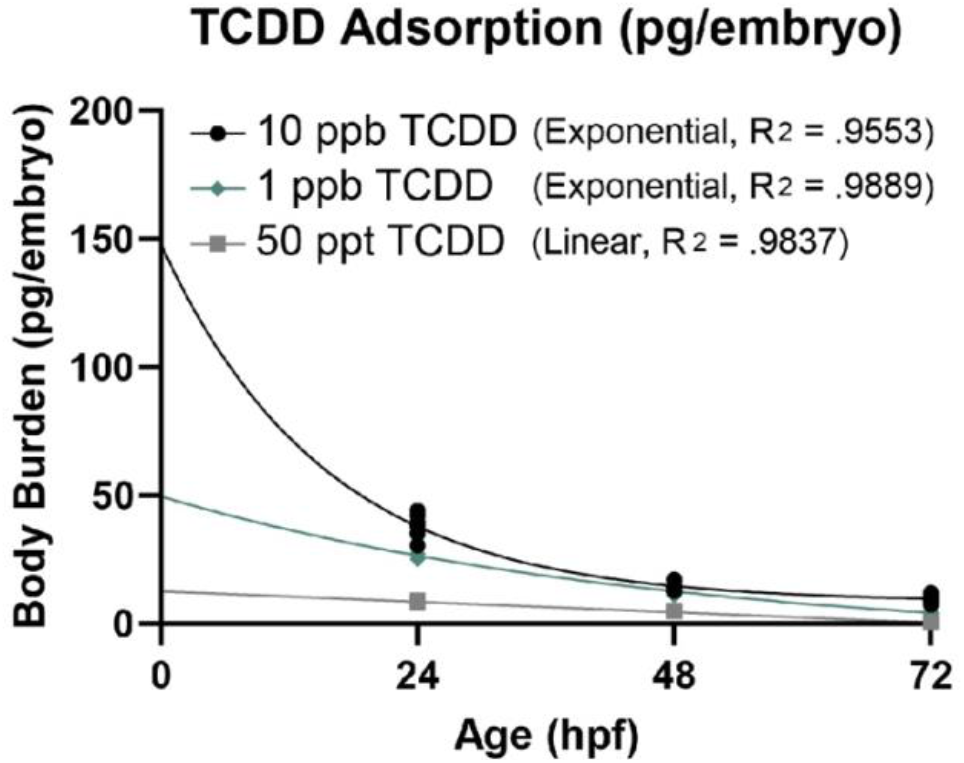
Embryonic TCDD concentration over time. Zebrafish were exposed to 10 ppb (black circles), 1 ppb (blue diamonds), and 50 ppt (grey boxes) at 4 hours post-fertilization (hpf). Samples were fixed at 24, 48, and 72 hpf and TCDD was quantified. 10 ppb and 1 ppb are modeled by exponential decay while 50 ppt uses linear elimination.

As expected, TCDD was not detected in any of the paired control samples at any of the timepoints examined. We found embryos dosed with 50 ppt TCDD for one hour resulted in a body burden of 8.53 ± 0.341 pg/embryo at 24 hpf (Table 1). We measured TCDD in a set of embryos exposed to 1 ppb TCDD and 10 ppb and, interestingly, found that the body burden of TCDD in the 1 ppb samples was similar to the 10 ppb samples. The 1 ppb does was determined to be 26.6 ± 1.21 pg/embryo and the 10 ppb dose, 38 ± 4.34 pg/embryo. At 24 and 72 hpf, there was a significant difference in TCDD body burden in the 1 ppb and 10 ppb dose, however at 48 hpf the body burden was not statistically different. This suggests that the body burden of TCDD at 24 hpf is not directly proportional to the exposure concentration at 4 hpf. A more detailed pharmacokinetics study of the zebrafish embryo is needed to dissect this proportional difference.

**Table 1.**
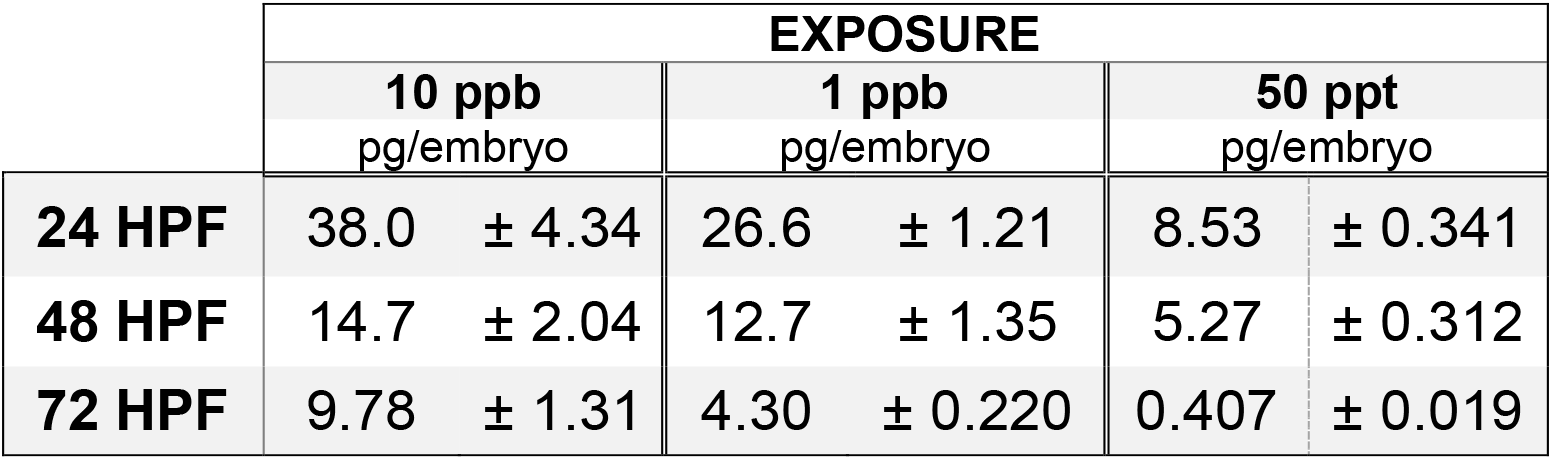
Concentration of TCDD detected per embryo over time. All values are mean ± standard deviation.

|  | EXPOSURE |  |  |  |  |  |
| --- | --- | --- | --- | --- | --- | --- |
|  | 10 ppb<br>pg/embryo |  | 1 ppb<br>pg/embryo |  | 50 ppt<br>pg/embryo |  |
| 24 HPF | 38.0 | ± 4.34 | 26.6 | ± 1.21 | 8.53 | ± 0.341 |
| 48 HPF | 14.7 | ± 2.04 | 12.7 | ± 1.35 | 5.27 | ± 0.312 |
| 72 HPF | 9.78 | ± 1.31 | 4.30 | ± 0.220 | 0.407 | ± 0.019 |

To contextualize zebrafish as a model of human health and ecotoxicology, we compared our findings to blood serum concentrations of TCDD detected in exposed human populations. From 1962 to 1971, American Vietnam veterans serving in Operation Ranch Hand in the Vietnam War sprayed Agent Orange, a defoliant contaminated with TCDD. As a result, serum dioxin concentration of TCDD in Operation Ranch Hand veterans have been shown to be high, measuring 12.2 ppt (51). However, the Vietnamese population endured significantly higher TCDD exposure. In 1970, human breast milk contained up to 1,832 ppt TCDD (excluding other dioxin congeners), by 1991 serum TCDD concentrations were found to be up to 33 ppt in the southern Vietnamese population, where Agent Orange was most heavily applied (52). In 1978, an industrial accident outside of Seveso, Italy, exposed residents to high concentrations of TCDD. This resulted in serum concentrations up to 56,000 ppt (56 ppb) in this population (53). Outside of large scale exposure events, the primary route of dioxin exposure is through dietary consumption of animal products, which results in *in utero* exposure and continuous exposure across the lifespan (54). Schecter et al. (54) determined the toxic equivalency levels (TEQ) in human food sources for dioxin and dioxin-like compounds. The TEQ is the amount of TCDD required to produce the same level of AHR activation as produced by the diverse composition of AHR agonists seen in environmental samples (55). Freshwater fish contained the highest TEQ (1.72 ppt), while a purely vegan diet had the lowest TEQ (0.09 ppt), intermediate levels were detected in chicken (0.33 ppt), ocean fish (0.39 ppt), and human milk (0.42 ppt, (54)). Using the previously reported dechorionated dry weight of a zebrafish embryo (39.5 ± 15.5 μg) at 24 hpf, we calculated the TCDD concentration within each embryo in our experiments and found that following: the 10 ppb exposure the internal TCDD concentration was approximately 0.96 ppt, 1 ppb exposure had an internal TCDD concentration of 0.67 ppt, and a 50 ppt dose the internal concentration was 0.22 ppt (56). This rough estimate suggests that our measured tissue concentrations of TCDD were within the same order of magnitude observed in human food sources and reflect environmental relevant levels of TCDD.

The described 10 ppb exposure paradigm produces an internal body burden that reflects the body burdens of TCDD in range of wild fish populations. The EPA reported TCDD concentrations in fish (all encompassing) to be about 6.89 ppt in 1999 (57). More recently, in 2008, the state of Maine reported TCDD concentrations to be between 1 and 2 ppt in freshwater fish. Although this concentration is significantly lower than the EPA report from 20 years ago, the 1-2 ppt concentration still poses a threat to human health and precautions related to fish consumption should be observed (58).

There are a number of variables to consider when comparing the body burden achieved from the current laboratory dosing paradigms and the body burdens that result from environmental exposures in wild fish populations. For example, a previous study found that repeated doses of 0.2 ppb or 50 ppt TCDD at 4.8 hpf and 31.2 hpf resulted in significant bioaccumulation of TCDD in embryonic zebrafish, producing a TCDD body burden between approximately 1 ppb and 5 ppb (59). The TCDD body burdens in wild fish populations, therefore, likely reflect repeated low-level exposures, which may produce different physiological effects than a single high dose exposure during early embryogenesis. Maternal body burden of TCDD is another important variable to model as it contributes to the earliest developmental events occurring at the time of fertilization. Female zebrafish dosed with TCDD provide a maternal load of TCDD in their eggs which is linearly portioned to adult body burden (60). Together, these findings highlight that different dosing paradigms result in different body burdens and emphasize the need to quantify body burden after varying aquatic exposures in order to make cross-study comparisons.

To determine if elimination of TCDD from the embryo occurred in a dose dependent manner, we modeled TCDD elimination for each dose using linear and exponential regressions. In our study, “elimination” captures all forms of removal of the parent compound, TCDD, including metabolism and excretion. Using a Comparison of Fits (Prism^®^), the 50 ppt exposed group could be fit by both an exponential (R^2^=0.983) and linear regression (R^2^=0.983); however, exponential decay could not accurately predict a half-life at 50 ppt. Therefore, the most parsimonious explanation, a linear regression (Elimination rate = 0.17 pg/hour), was used to represent the 50 ppt exposure group. The 1 ppb and 10 ppb group were best fit by exponential decay (R^2^=0.9889 and 0.955, respectively) compared to a linear regression (R^2^=0.9650 and 0.846, respectively). Using first-order kinetics, we calculated the half-life of TCDD in the zebrafish embryos following a 1 ppb exposure to be 32.71 hours (95% Confidence Interval (CI) = 19.34 – 85.36 hours), while the 10 ppb dose had a half-life of 10.72 hours (95% CI = 7.336 - 19.88 hours). Previous studies in humans demonstrate that elimination of TCDD occurs through first order kinetics, meaning that the relative concentration of parent compound determines the rate of elimination (61). It is possible that if we measure TCDD concentration prior to 24 hpf in the 50 ppt dose, the data would be more representative of first order kinetics.

### Metabolic pathway analysis

After establishing that the 10 ppb or 50 ppt embryo dosing paradigm used in zebrafish to study developmental toxicology is representative of relevant body burdens found in human exposure and wildlife populations, we next used non-targeted LC-HRMS to evaluate metabolic changes of embryonic zebrafish that result from TCDD exposure. Metabolic changes were assessed at 24, 48, and 72 hpf for both the highest, 10 ppb, and lowest, 50 ppt, doses of TCDD. Reverse phase chromatography was used for negative ionization, which separates compounds with hydrophobic moieties. Normal phase chromatography was used for positive ionization, which differentiates compounds with hydrophilic moieties. A total of 6,874 m/z features were detected in positive ionization mode, while 5,565 m/z features were detected in negative ionization mode for all exposure levels and time points. We used all features detected for significance testing.

We performed metabolomics using t-tests for each exposure dose and time point relative to the control samples. We first performed metabolomics analysis on the 50 ppt and 10 ppb exposure groups over the three time points in negative ion mode (Figure 2, Table S1) and then repeated this analysis in positive ion mode (Figure 3, Table S1). In the 50 ppt group,10 features were significant in negative ionization mode and 2 features were significant in positive ion mode. In the 10 ppb exposure group, 1,224 features were significant in positive ionization mode and 870 features were significant in negative ionization mode. The features that were significant in the 50 ppt dosing scheme were not significant in the 10 ppb dosing scheme. These findings indicate that the lower 50 ppt dose TCDD activates different metabolic processes than a high dose. Future study on this topic is needed to completely understand this phenomenon.

**Figure 2.**
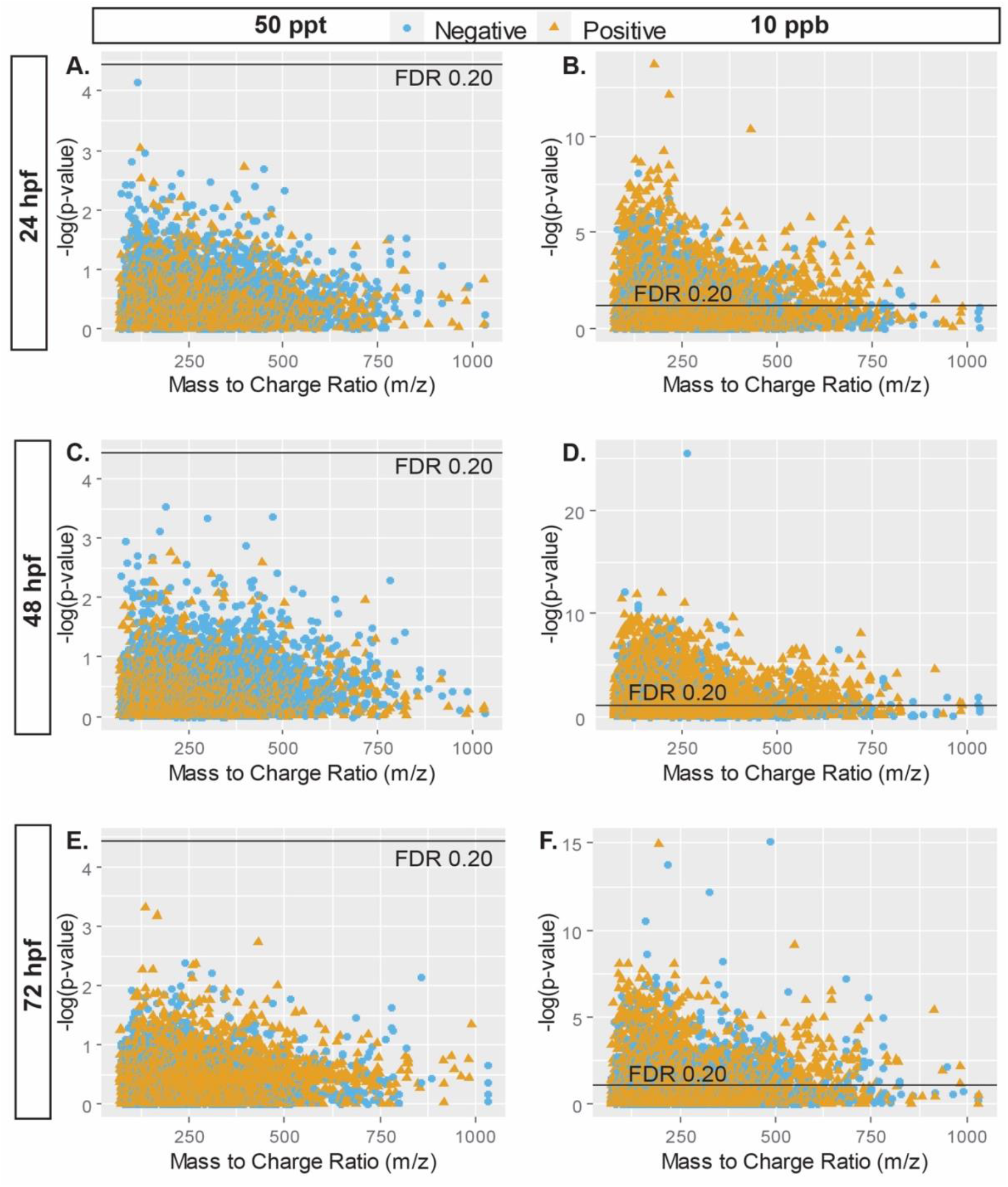
Manhattan plots of log 10 transformed p-values for the association between m/z feature and 50 ppt (A – 24 hpf, C – 48 hpf, and E – 72 hpf) and 10 ppb (B – 24 hpf, D – 48 hpf, and F – 72 hpf) TCDD exposure in negative ionization mode. Blue circles represent features that were down regulated in comparison to the control group. Orange triangles represent features that were up regulated in comparison to the control group. The black line is the p-value corresponding to the false discovery rate (FDR) threshold of 0.2.

**Figure 3.**
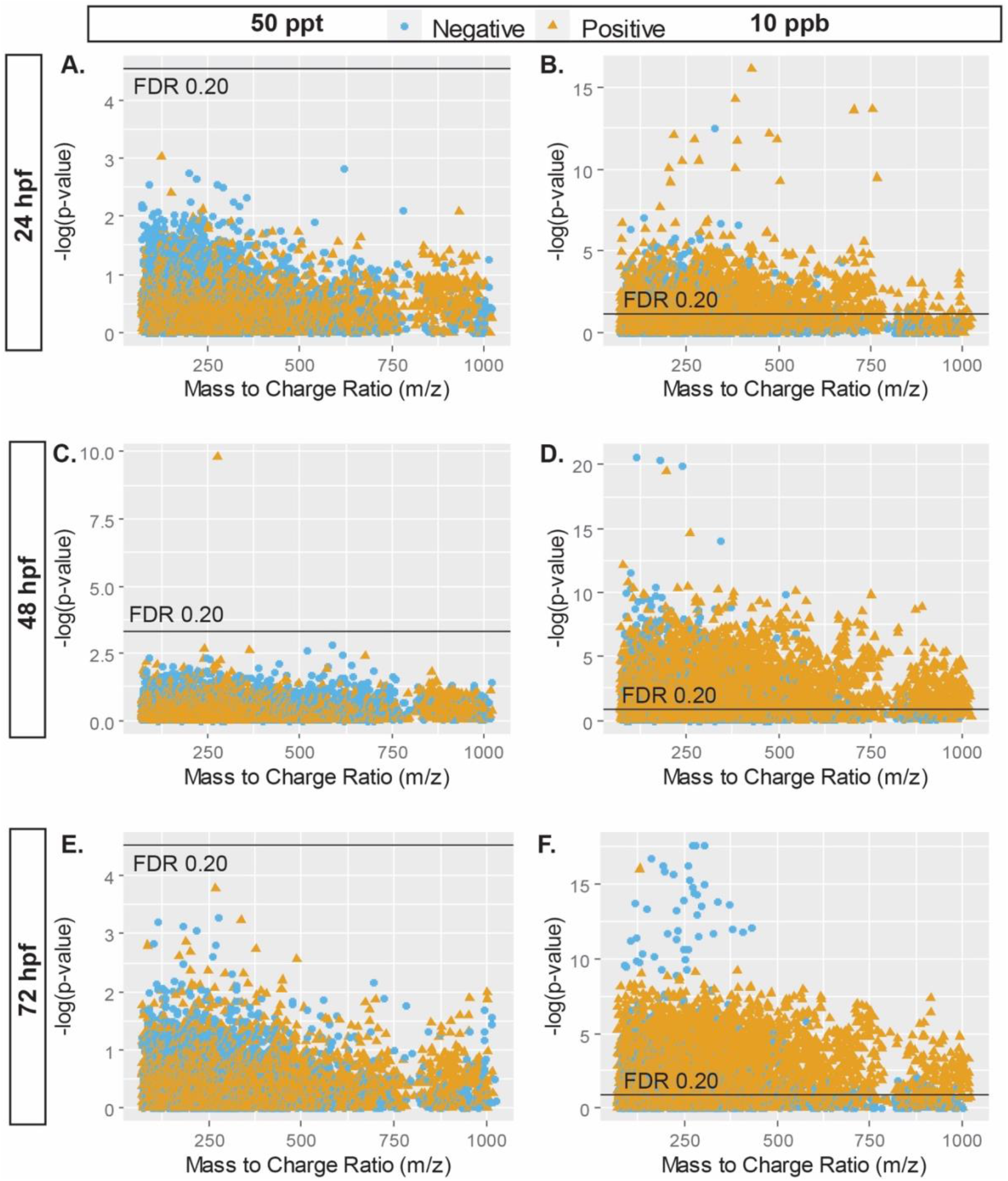
Manhattan plots of log10 transformed p-values for the association between m/z feature and 50 ppt (A – 24 hpf, C – 48 hpf, and E – 72 hpf) and 10 ppb (B – 24 hpf, D – 48 hpf, and F – 72 hpf) TCDD exposure in positive ionization mode. Blue circles represent features that were down regulated in comparison to the control group. Orange triangles represent features that were up regulated in comparison to the control group. The black line is the p-value corresponding to the false discovery rate (FDR) threshold of 0.2.

Given that there were a small number of significant features detected in the 50 ppt exposure group, pathway enrichment analysis was only performed on the 10 ppb exposure group and the associated pathways are shown in Figure 4. Hit totals and significance can be found in Table S2. In total, 85 pathways were enriched: 23 pathways enriched in both modes, 1 pathway (Glycosylphosphatidylinositol(GPI)-anchor biosynthesis) unique to the positive ionization mode, and 61 pathways unique to the negative ionization mode (Figure 4, Table S2).

**Figure 4.**
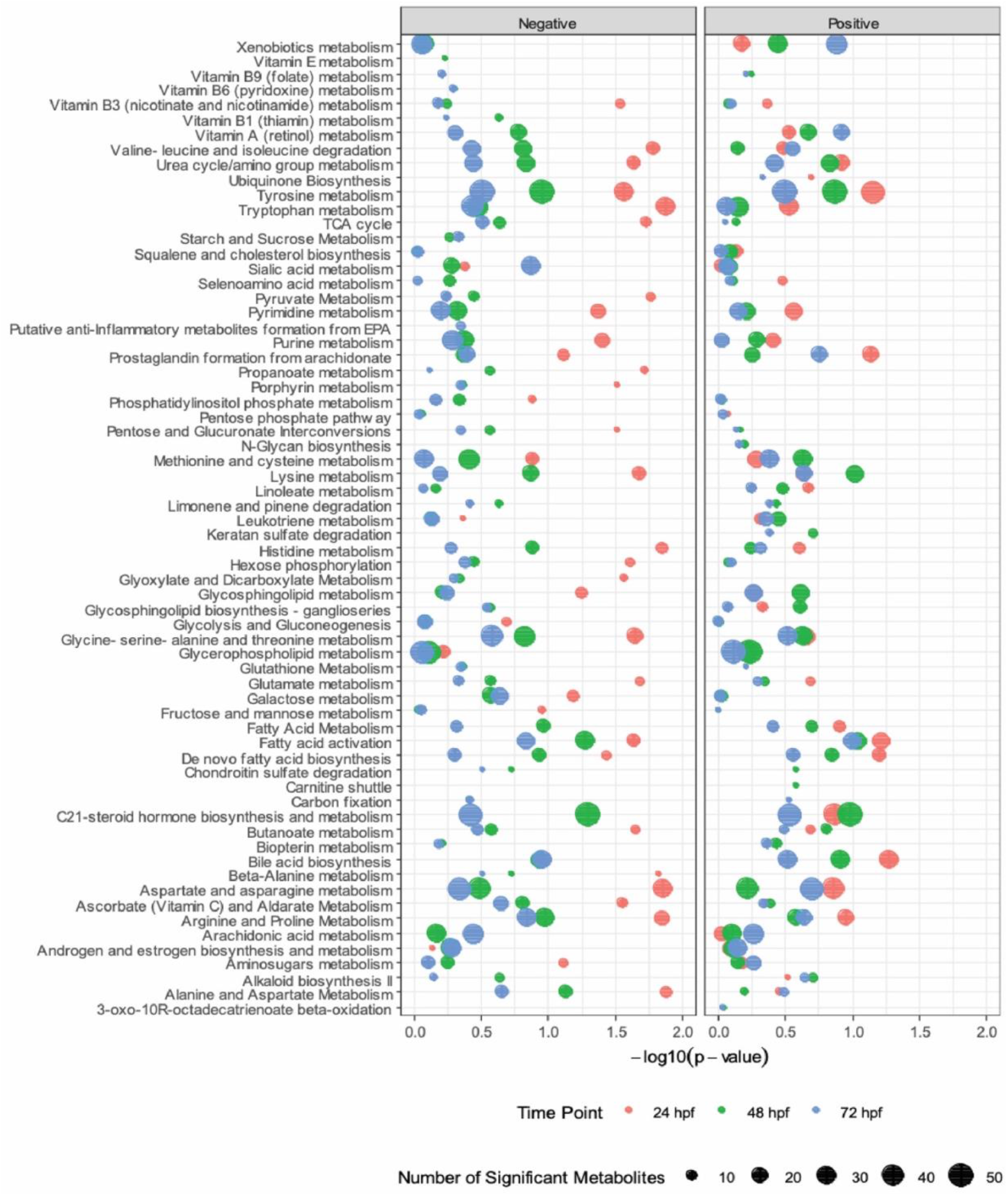
Enriched pathways associated with 10 ppb TCDD exposure in negative and positive ionization mode with at least 5 significant metabolites. The size of the dot represents the number of overlapping metabolites in the pathway. The position of the dot is determined by the −log p-value obtained from 10,000 permutations.

### Metabolomic results accurately represent known effects of TCDD exposure

We used previously published endpoints associated with TCDD exposure to authenticate our metabolomics results. The metabolism of TCDD has been well characterized; however, we were unable to detect the direct metabolites of TCDD that have been previously described in mammals (62–64). This is likely because these studies were performed on bile or liver cell excretions, isolated liver tissue, or cultured hepatocytes (62–64). Therefore, these metabolites would have been enriched in these samples compared to the proportionally lower level of abundance in our pooled whole embryo homogenates. Nonetheless, the xenobiotic metabolism pathway was associated with TCDD exposure (Figure 3). Xenobiotic metabolism is a generalized category of metabolites that are produced after exposure to an exogenous chemical. This pathway includes the breakdown of aromatic hydrocarbons and is highly associated with cytochrome P450 induction. Dioxins, all of which are aromatic hydrocarbons, activate expression of cytochrome P450s in zebrafish. Appropriately, *cyp1a* is used as a biomarker of dioxin exposure (44). The most significant pathways dysregulation was observed at 24 hpf (higher −log10(P)). For most pathways, the effects were mitigated over time (lower - log10(P)); however, with the exception of xenobiotic metabolism pathway, which became more dysregulated over time (Figure 4). The upregulation of metabolites in this pathway over time suggests that the elimination of TCDD observed between 24 and 72 hpf could be due to metabolism of aromatic TCDD rather than excretion (Figure 1).

Our confidence in the metabolomics findings is further supported by the observed changes in the tryptophan metabolism pathway. Tryptophan has been identified as the endogenous AHR ligand in both mammals and zebrafish and, in rats, exposure to TCDD causes a dose-related change in tryptophan metabolism (65–67). Consistent with these previous reports, we found tryptophan metabolism pathway was associated with TCDD exposure in zebrafish; thus, indicating that exogenous AHR activation by TCDD may affect the natural biosynthesis of tryptophan and contribute to the pathophysiology associated with TCDD exposure. The association of the tryptophan metabolism pathway with TCDD exposure supports previously characterized effects of TCDD exposure on pathways initiated by AHR activation.

Bile acid metabolism was another significantly altered metabolic pathway associated with TCDD exposure. Recent work in mice has shown that bile acid biosynthesis is altered after TCDD exposure; however, this change is only detected in a liver biopsy and not detected in the serum (68). Thus, the only way to observe these effects is through more invasive methods in mammals. Using zebrafish embryos, we are able to detect changes to the metabolome which may be masked through less invasive methods of discovery in mammals or humans, such as blood or urine collection. This result emphasizes the utility of the zebrafish embryo to detect metabolic alterations that signify toxic effects.

A total of 224 unique m/z features were identified in the enriched metabolic pathways. Of these features, 80 were up regulated features and 144 were downregulated (Table S2). The feature annotated as dihydrotestosterone (DHT) was one of the most significantly downregulated metabolites after TCDD exposure. DHT peak intensity decreased 6.02-fold every 24 hours. DHT is a major component of the androgen and estrogen biosynthesis and metabolism pathway, as is the interconnected C21-steroid hormone biosynthesis and metabolism pathway. It is well established that exposure to TCDD effects endocrine signaling and reproductive endpoints in humans, mammals, and zebrafish (36,46,69–78). In a study of male infertility, Galimova et al. (79) found that infertile men had significantly higher concentrations of dioxin in their semen compared to fertile men. Additionally, in two different studies of human populations exposed to high concentrations of environmental dioxin, both adult men and children had altered levels of DHT compared to control populations (80,81). Thus, our metabolomic analysis in zebrafish embryos after TCDD exposure reflect what is seen in a subset of human exposure conditions. This not only validates our finding, but also, supports the conclusion that zebrafish can be used to model metabolic changes related to hormonal pathways in humans.

### TCDD exposure induces changes in metabolic pathways important for brain development

TCDD is a highly toxic environmental contaminant known to target many different biological systems including the brain. Our metabolomic analysis revealed dysregulation of critical brain metabolic pathways, including glutamate metabolism, tyrosine metabolism, and chondroitin sulfate degradation. TCDD exposure altered glutamate metabolism in the zebrafish embryo. In this pathway, gamma-aminobutyric acid (GABA) was downregulated 0.32-fold over time and L-Glutamic acid (glutamate) was downregulated 0.30-fold over time. GABA, as well as glutamate are key neurotransmitters in the brain that are essential for establishing the balance of excitation and inhibition in the brain (82–85). Dysregulation of excitation and inhibition is implicated in a broad range of neurodevelopmental and neuropsychiatric diseases including epilepsy, autism, ADHD, and schizophrenia ((86–89), as reviewed in (90). Previous studies using hippocampal slices and xenopus oocyte preps have shown that GABAergic signaling is significantly affected by non-dioxin-like PCBs (91,92). The non-dioxin-like PCBs are nonplanar and have one or more chlorines in the *ortho*-positions, whereas dioxin-like TCDD has a planar (flat) structure. Our results suggest that TCDD may also alter GABAergic signaling and excitatory/inhibitory balance in the developing brain, a topic currently being investigated by our group.

TCDD exposure was associated with altered metabolism of multiple neurotransmitters. Neurotransmitters are signaling molecules released by neurons to communicate and stimulate other cells including neurons, oligodendrocytes, astrocytes, microglia, and muscles. Tyrosine biosynthesis was significantly associated with TCDD exposure and resulted in a 0.66-fold downregulation of dopamine over time. Dopamine is a neurotransmitter that is fundamental component of the reward system and is also critical for the initiation and coordination of motor sequences (93,94). In mammals, dopamine is downregulated in the brain after exposure to TCDD likely through the inhibition of tyrosine hydroxylase (95). It has been previously shown that serum dioxin concentrations are associated with the development of attention deficit hyperactivity disorder (ADHD, (96)). In negative ion mode 19 features were associated with TCDD exposure (p=0.089), and in positive ion mode 2 features compounds were significantly associated with exposure (p= 0.012). This suggests that TCDD may not only alter dopamine biosynthesis, but also the formation of other neurotransmitters essential for brain function. Consequently, our results support that TCDD exposure and other AHR agonists could contribute to the formation of other disease states caused by changes in dopamine signaling such as Parkinson’s Disease, and multiple sclerosis (97,98).

We hypothesize there are two potential reasons chondroitin sulfate degradation was altered by dioxin exposure. First, chondroitin sulfate proteoglycan (CSPG) biosynthesis is involved in cartilage formation (99). It is well known that TCDD exposure causes craniofacial defects and prior studies demonstrate TCDD exposure in zebrafish causes decreased chondrocytes number within the jaw cartilage of the developing bone (37,100,101). Therefore, it is possible that chondroitin sulfate degradation was altered due to the reduced number of chondrocytes. However, CSPG biosynthesis is also an essential part of the extracellular matrix (ECM) formation of the brain (102). In mammal and zebrafish, TCDD exposure is known to cross the BBB and alter neural and glia cell number and function (49,103). During brain development, CSPG bind signaling molecules, such as fibroblast growth factors, to affect intercellular signaling (104). Consequently, it is also possible that TCDD targets the brain by altering the formation of this essential ECM component. To the best of our knowledge, TCDD induced changes in ECM composition in the brain have not been previously reported.

### Conclusions

Together, our findings provide fundamental information that facilitates the use of zebrafish as a model for understanding TCDD-induced toxicity. We employed targeted GC-HRMS analysis to quantify TCDD concentrations in the zebrafish embryo at key developmental timepoints. We found that TCDD is eliminated by exponential decay after a 1 ppb and 10 ppb exposure, while after a lower, 50 ppt dose, a linear model better captures TCDD elimination. Embryonic and larval body burdens of TCDD for both doses were previously unknown. Therefore, this study provides essential information that helps contextualize studies using this common dosing paradigm. We used non-targeted high-resolution LC-HRMS to detect metabolites in the zebrafish embryo and then performed metabolomic pathway analysis to identify known metabolic pathways that were affected by TCDD exposure. Zebrafish continue to be a powerful model for studying the health effects of dioxin exposure as metabolomics revealed pathways that are not detectable in serum, blood or urine samples, such as bile acid biosynthesis. Further, neurotransmitter pathways were associated with TCDD exposure, demonstrating that the study of the zebrafish metabolome after TCDD exposure may reveal previously overlooked effects of dioxin exposure on the health of humans and aquatic species environment.

## ABBREVIATIONS

AHR: Aryl-Hydrocarbon Receptor
CSPG: Chondroitin sulfate proteoglycan
DHT: Dihydrotestosterone
ECM: Extracellular Matrix
FDR: False Discovery Rate
GABA: Gamma-Aminobutyric acid
GC: Gas chromatography
hpf: Hours post-fertilization
HRMS: High resolution mass spectrometry
LC: Liquid chromatography
PCB: Polychlorinated Biphenyl
ppb: parts per billion
ppt: parts per trillion
TCDD: 2,3,7,8-Tetrachlorodibenzo-p-dioxin

## ACKNOWLEDGMENTS

Thank you to Jessica Plavicki and Kurt Pennell for their encouragement and support of this COVID inspired project which was executed through a series of hand offs and Zoom meetings.

## SUPPORTING INFORMATION

**Table S1.** Significant features detected in positive ion mode **(a)** and negative ion mode **(b)** after 50 ppt exposure. Significant features detected in positive ion mode **(c)**, and negative ion mode **(d)** after 10 ppb exposure.

**Table S2.** Pathway total features and significance after 10 ppb exposure in positive ion mode **(a)**, and negative ion mode **(b).** Annotations of the significant features detected in pathway analysis and their regulation in positive ion mode **(c)**, and negative ion mode **(d)**.

**Figure S1.** Stock Solution and working solution concentrations validated with GC-HRMS.

