## Supplementary material for "High-resolution mass spectrometry reveals environmentally relevant uptake, elimination, and metabolic alterations following early embryonic exposure to 2,3,7,8-tetrachlorodibenzo-p-dioxin in zebrafish": Figure S1

| **Stocks** | **Expected Concentration** | **Actual Concentration (mean±SD)** |
| --- | --- | --- |
| Stock A | 50,000 ppb | 50.00 ± 0.32* µg/mL (ppm) |
| Stock B | 10,000 ppb | 10,280[†] ng/mL (ppb) |
| Stock C | 1,000 ppb | 1,003.869[†] ng/mL (ppb) |
| Stock D | 50,000 ppt | 50.411 ± 0.274[‡] ng/ml (ppb) |
| **Working solution** | **Expected Concentration** | **Actual Concentration (mean±SD)** |
| Diluted from Stock B | 10 ppb | 9.973 ± 0.223[‡] ng/mL (ppb) |
| Diluted from Stock C | 1 ppb | 1.045 ± 0.197[‡] ng/mL (ppb) |
| Diluted from stock D | 50 ppt | 50.13 ± 0.98[‡] pg/mL (ppt) |

* Measured by the manufacturer
[†] Measured in technical triplicate
[‡] n=3

**Figure S1.** Stock Solution and working solution concentrations validated with GC-HRMS.
